# Region Specific Central Arbor Morphologies of Nociceptive Afferents Develop Independently of Their Peripheral Target Innervation

**DOI:** 10.1101/285098

**Authors:** William Olson, Wenqin Luo

## Abstract

Functionally important regions of sensory maps are overrepresented in the sensory pathways and cortex, but the underlying developmental mechanisms are not clear. In the spinal cord dorsal horn (DH), we recently showed that paw innervating Mrgprd+ non-peptidergic nociceptors display distinctive central arbor morphologies that well correlate with increased synapse transmission efficiency and heightened sensitivity of distal limb skin. Given that peripheral and central arbor formation of Mrgprd+ neurons co-occurs around the time of birth, we tested whether peripheral cues from different skin areas and/or postnatal reorganization mechanisms could instruct this somatotopic difference among central arbors. We found that, while terminal outgrowth/refinement occurs during early postnatal development in both the skin and the DH, postnatal refinement of central terminals precedes that of peripheral terminals. Further, we used single-cell ablation of Ret to genetically disrupt epidermal innervation of Mrgprd+ neurons and revealed that the somatotopic difference among their central arbors was unaffected by this manipulation. Finally, we saw that region-specific Mrgprd+ central terminal arbors are present from the earliest postnatal stages, before skin terminals are evident. Together, our data indicate that region-specific organization of Mrgprd+ neuron central arbors develops independently of peripheral target innervation and is present shortly after initial central terminal formation, suggesting that either cell-intrinsic and/or DH local signaling may establish this somatotopic difference.

## INTRODUCTION

Animals use distal limb regions (paws/hands) for exploration and object manipulation. To facilitate these behavioral requirements, distal limb somatosensory circuits have several region-specific (somatotopic) organization mechanisms to increase the sensitivity of these skin regions. One mechanism involves differences in primary neuron density in the periphery [1, 2]. In addition, distal limb representations could be ‘magnified’ through region specific circuit organization in the central nervous system, leading to its overrepresentation in the spinal cord and cortex circuits [3-5]. While other sensory systems use analogous forms of regional magnification in various species [6-9], the developmental mechanisms used by sensory systems to differentially allocate circuit space in this manner have not been clearly defined.

Similar to light touch, humans have increased sensitivity for pain in the distal limbs and fingertips [10, 11]. Recently, we used single-cell genetic tracing of Mrgprd+ mouse nociceptors to characterize the somatotopic organization of mammalian pain neurons [12]. While we found no obvious peripheral mechanisms for increased sensitivity in the paw (these neurons have lower innervation density in the paw glabrous skin compared to the limb hairy skin and have similar terminal areas across skin regions), we identified a novel region-specific organization of the central terminal arbors of these neurons in the spinal cord dorsal horn (DH) (Figure 1A). Specifically, paw and trunk innervating nociceptors have distinct terminal morphologies (‘round’ vs. ‘long’) in the DH, such that paw nociceptors have a much wider mediolateral spread. Interestingly, this ‘round’ central terminal arbor morphology closely correlates with increased synaptic transmission efficiency in the spinal cord and decreased threshold for activating these afferents in the paw skin. This work suggests that ‘magnification’ of these nociceptors through region-specific central arbors could expand the representation of the paw in the spinal cord and downstream circuits. These results also indicate that Mrgprd+ neurons, which as a population show very similar molecular markers and gross anatomical features, differentially direct their central terminal arbor formation based on their somatotopic location during development. Our ability to trace and genetically manipulate single primary afferents from this population offers a unique ability to study the developmental mechanisms underlying region-specific organization of this sensory circuit.

**Figure 1.**
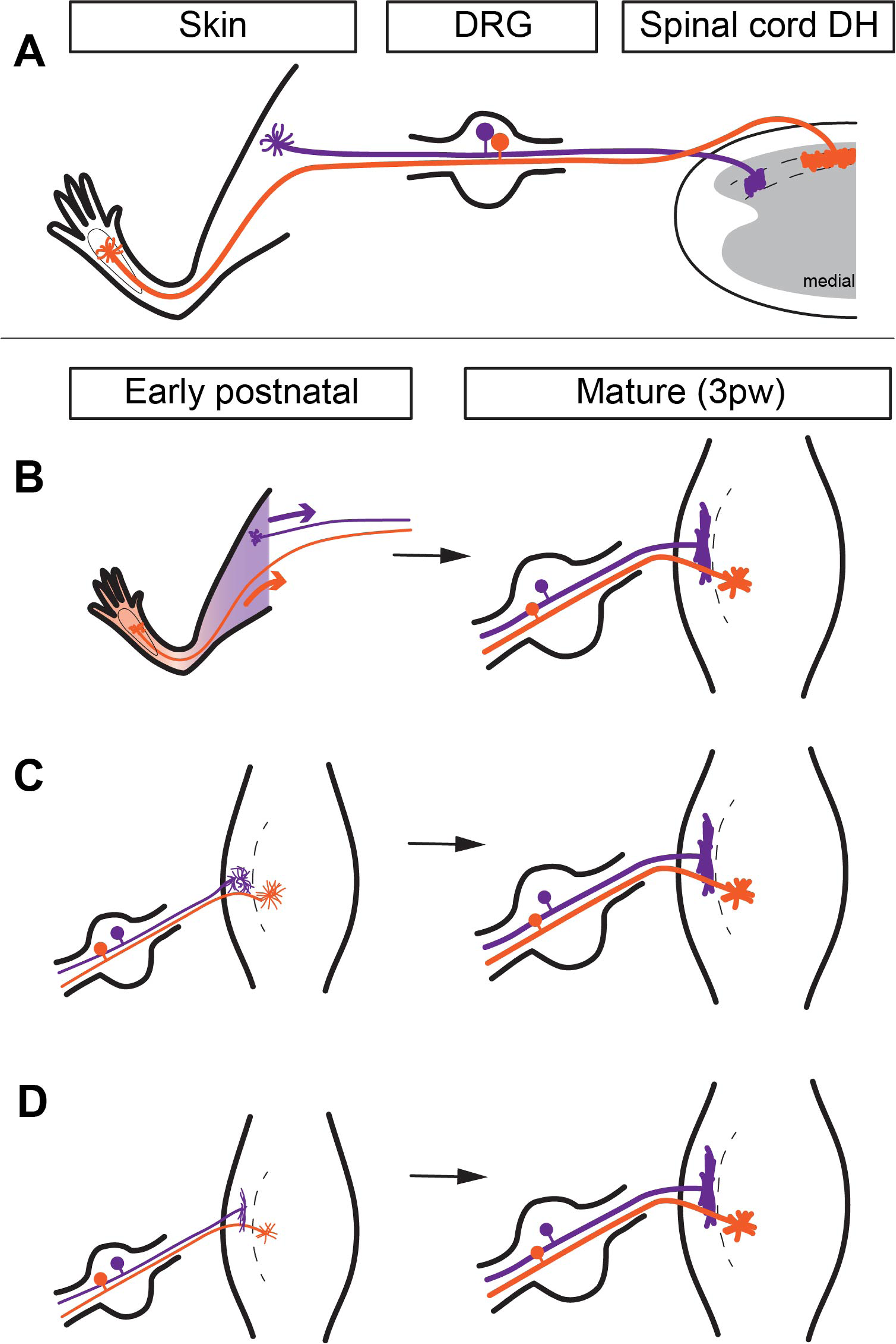
Models of hypothesized developmental mechanisms for region-specific Mrgprd+ central arbors. A, Mature somatotopic organization of Mrgprd+ afferents. Proximal hind limb afferents (purple) grow ‘long and thin’ central arbors in the lateral DH, while plantar paw afferents (orange) grow ‘round and wide’ central arbors in the medial DH. DH is drawn as a transverse section. B, Peripheral cues model. Upon innervation of the skin around birth, cues from different skin regions direct region-specific development of central terminal arbors. DH is drawn from a top-down view. C, Central reorganization model. Afferents grow immature homogenous arbors that are postnatally reorganized into region-specific arbor morphologies. D, Pre-patterned model. Afferents grow somatotopically distinct arbor morphologies during their initial innervation of the DH.

The mechanisms that direct somatotopically-appropriate wiring of DRG neurons are largely unclear. Somatosensory circuits of the DH have a ‘flipped’ topographic map: in the lumbar enlargement (innervating the hindlimbs), the distal limbs (foot and toes) are represented in medial DH while the proximal limbs are represented in the lateral DH (Figure 1A) [13, 14]. Nerve tracing experiments have shown that cutaneous sensory topographic innervation maps formed early in development are similar to the mature pattern [15, 16]. Based on the rough coincidence of peripheral and central target innervation, and based on the proximal-to-distal progression of hind limb epidermal innervation, it was proposed that peripheral innervation could drive correct topographic innervation pattern in the DH [17]. However, subsequent studies suggested that peripheral and central topographic innervation may develop independently of one another [18-20].

Despite the interesting information gained from these experiments, the previous work did not resolve the single-cell structure of DRG neurons. Therefore, these studies could not examine the mechanisms underlying the disproportionate representation (magnification) of paw regions in somatosensory circuits. It remains possible that peripheral cues from different skin regions could instruct the formation of region-specific central arbor morphologies of Mrgrd+ neurons in the DH (Figure 1B). Alternatively, it is also possible that Mrgrd+ neurons form immature homogenous arbors that are postnatally reorganized into region-specific morphologies (Figure 1C). Lastly, DRG afferents may form region-specific arbor morphologies during their initial terminal formation, suggesting pre-patterning mechanisms (Figure 1D). Here, we used population-level tracing to characterize the postnatal development of Mrgprd+ nociceptor central and peripheral terminal arbors. In addition, we performed single-cell ablation of Ret to disrupt peripheral target innervation of these neurons and analyze the effect on their central arbor morphology in the DH. Lastly, we performed single-cell tracing of Mrgprd+ neurons in early postnatal animals, right after their initial innervation of the DH. These experiments show that region-specific arbors are present in early postnatal animals (supporting the ‘pre-patterned’ model), and that central terminal development slightly precedes, and occurs independently of, peripheral terminal development/refinement. Taken together, our results suggest that somatotopic organization of mammalian nociceptor central terminal arbors is likely to be dictated through mechanisms intrinsic to the DRG neurons themselves and/or by mechanisms within the spinal cord.

## RESULTS AND DISCUSSION

### Population-level characterization of postnatal development of peripheral and central terminals of Mrgprd+ DRG neurons

To investigate whether region-specific arbor development may be driven by peripheral and/or central mechanisms, we used *Mrgprd*^*EGFPf*^ knock-in mice [21] to characterize the postnatal (P1 – 3 postnatal weeks, pw) innervation of non-peptidergic nociceptors in the paw glabrous skin and the lumbar spinal cord enlargement DH. *Mrgprd* is first expressed at E16.5 in mice and specifically marks the non-peptidergic nociceptor population [22]. EGFP expression in this knock-in mouse line faithfully indicates expression of *Mrgprd*. This genetic tool offers advantages over previous approaches since it specifically labels non-peptidergic fibers (unlike nerve filling [18, 23]) and avoids issues related to the dynamic expression of immunostaining markers [17, 24].

Peripherally, mature non-peptidergic nociceptor axons travel to the skin in the cutaneous nerves, grow a fiber plexus parallel to the skin surface in the dermis, and send perpendicular terminals out of the subepidermal plexus that penetrate the epidermis [12, 21]. Most of the paw epidermis is not innervated by *Mrgprd*^*EGFPf*^ fibers at P1, except for few rudimentary terminals (Figure 2A). From P1 to P7, there is a rapid phase of nerve terminal growth, as indicated by a great increase in density of both primary (leaving the subepidermal plexus) as well as secondary/tertiary (branches off primary terminals) *Mrgprd*+ fiber branches (Figure 2A-C, K, M, N). It is then followed by a refinement phase, as indicated by a decrease in density from P7 to 3pw. By 3pw, no secondary or tertiary branches are present (Figure 2G, H, L-N). This pattern is true when quantified as absolute terminal densities or as growth-normalized values (Figure 2M, N).

Centrally, non-peptidergic nociceptor axons travel through the dorsal roots, grow for 0-2 spinal segments rostrally or caudally in Lisseur’s tract at the dorsolateral margin of the spinal cord, and then dive ventrally to innervate layer II of the DH [12, 21]. *Mrgprd*^*EGFPf*^ fibers have established a thick (in the dorsoventral extent) terminal layer by P1 in the DH (Figure 2D). This layer shows a 4-fold (absolute) decrease in layer thickness, reaching the mature layer thickness by P7 (Figure 2D-F, I, J). This decrease of layer thickness is also true when quantified by growth-normalized values (Figure 2O, P). In summary, while *Mrgprd*^*EGFPf*^ peripheral terminals are still undergoing initial outgrowth in the epidermis (the first postnatal week), their central terminals are in the process of refining to their mature thickness in the DH. These results indicate that central terminal development/refinement of *Mrgprd*+ neurons precedes peripheral development/refinement in the postnatal period.

**Figure 2.**
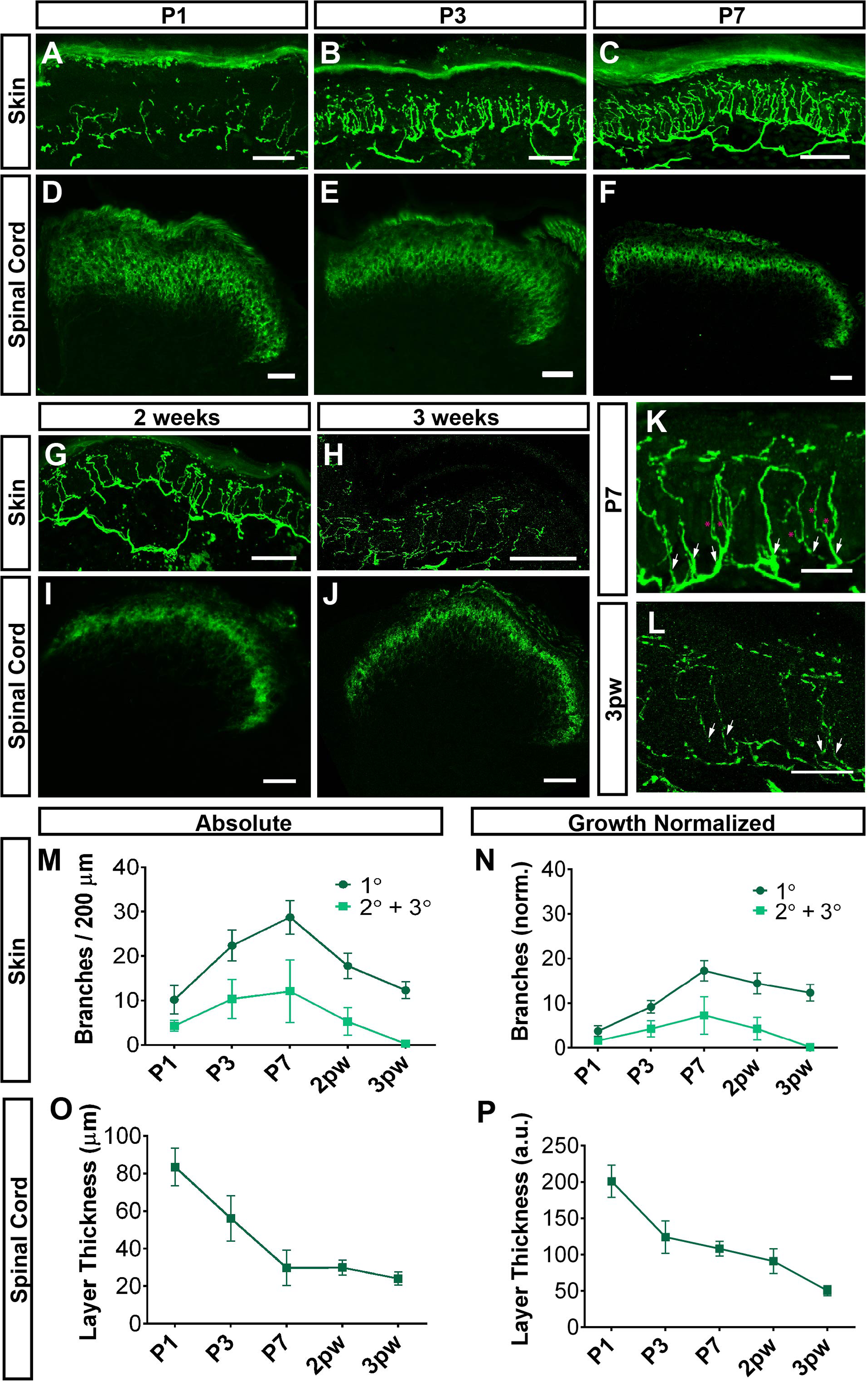
Postnatal central and peripheral terminal development of *Mrgprd*^*EGFPf*^ non-peptidergic nociceptors. A-J, GFP immunostaining of glabrous skin (A-C, G, H) and DH (D-F, I, J) sections from *Mrgprd*^*EGFPf*^ mice at the indicated ages. K, L, Higher magnification views of peripheral terminals, indicating secondary/tertiary branches (pink asterisks) growing off primary branches (white arrows) in P7 but not 3pw skin. M, N, Quantification of densities of absolute (M) and growth-normalized (N, see Methods) glabrous skin primary, secondary and tertiary branches during postnatal development. Skin terminals show overgrowth during the first week. O, P, Quantification of absolute (O) and growth-normalized (P, see Methods) DH layer thickness at the indicated ages. DH terminals show a refinement during the first week, at which point they remain at their mature thickness. *n* = 3 animals per stage. Scale bars = 50 µm (A-J), 20 µm (K, L).

### Non-peptidergic nociceptor central arbor formation is independent of peripheral terminal innervation

Next, we take advantage of a genetic manipulation to further determine whether non-peptidergic nociceptors utilize peripheral cues or processes to establish region-specific central arbor morphologies. Previous work has shown that, upon DRG-specific deletion of the Glial cell line-derived neurotrophic factor receptor Ret, non-peptidergic nociceptors fail to innervate their final peripheral target, the skin epidermis, while their central terminals remain in DH layer II [25]. However, these experiments did not trace single-cell morphology, so it remains unknown whether their central arbors are altered after this failure in peripheral target innervation. We ablated *Ret* from individual non-peptidergic nociceptors and quantitatively measured their regional single-cell width (mediolateral) in the DH. We used a mouse line in which Cre-dependent inactivation of Ret also leads to expression of CFP (*Ret*^*f(CFP)*^) [26]. In *Mrgprd*^*CreERT2 /+*^; *Ret*^*f(CFP)/ null*^ mice (Ret CKO), which carry the *Ret*^*f(CFP)*^ allele along with a Ret null allele, low-dose prenatal tamoxifen (0.5 mg at E16.5-E17.5) generated sparse Ret-null non-peptidergic nociceptors that are labeled with CFP (Figure 3C). In *Mrgprd*^*CreERT2 /+*^; *Ret* ^*f(CFP)/ +*^ littermate controls (Control), this same treatment sparsely labeled Ret heterozygous non-peptidergic neurons with CFP (Figure 3A). As expected for Ret deletion, sparsely labeled CFP DRG cell bodies were smaller in *Mrgprd*^*CreERT2 /+*^; *Ret*^*f(CFP)/ null*^ mutant mice (data not shown), but the number of CFP_+_ neurons was not decreased (average CFP_+_ neurons/DRG: control = 58.7 ± 10.3, n = 14 DRGs from 3 animals, mutant = 70.4 ± 24.6 n = 22 DRGs from 3 animals) (Figure 3A, C, I). In addition, while sections of glabrous skin from control mice showed sparsely labeled terminals with mature epidermal endings, sections from mutant skin showed axon bundles in the dermis but no mature endings in the epidermis, consistent with previous work [25] (Figure 3B, D). Lastly, serial sectioning of control and mutant thoracic (T6-T12) and lumbar (L3-L6) spinal cords showed that the round-vs.-long distinction in central terminal morphology was unaffected by Ret deletion (Figure 4E-H). We imaged through serial sections and measured the maximum mediolateral width of sparse-labeled neurons. The medial lumbar neurons were roughly twice as wide, on average, as either lateral lumbar or thoracic neurons (thoracic = 28.7 ± 5.2 μm, lateral lumbar = 33.5 ± 6.5 μm, medial lumbar = 64.7 ± 14.2 μm, n = 287 neurons from 3 mice) in control mice (Figure 3E&F, J). A similar difference was also seen in mutant spinal cords (thoracic = 29.0 ± 7.0 μm, lateral lumbar = 33.1 ± 8.7 μm, medial lumbar = 68.6 ± 18.1 μm, n = 239 neurons from 3 mice (Figure 3G&H, J), indicating that the round-vs.-long distinction is maintained in these mutant neurons. Additionally, no difference in the average width of round or long terminals was observed between control and Ret null non-peptidergic neurons (Figure 3J), suggesting that *Mrgprd*+ neurons grow normal central terminal arbor morphologies in the absence of Ret.

**Figure 3.**
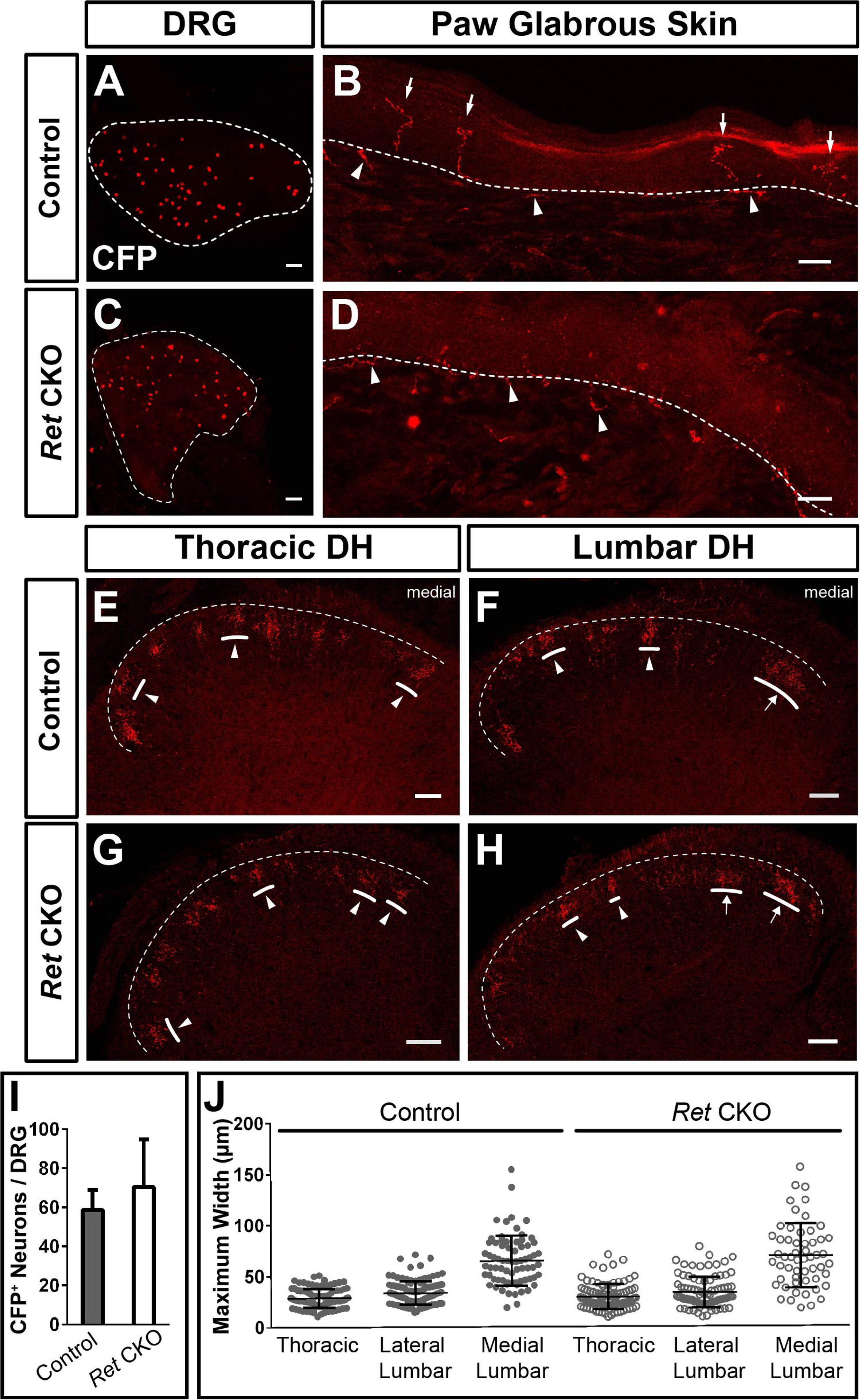
Genetic disruption of peripheral target innervation does not affect region-specific central arbor morphologies. A, C, Whole mount CFP immunostaining of sparse labeled 2-3pw *Mrgprd*^*CreERT2*^; *Ret* ^*f(CFP) / +*^ (control, A) and *Mrgprd*^*CreERT2*^; *Ret* ^*f(CFP) / null*^ (mutant, C) DRGs (0.5 mg tamoxifen at E16.5-E17.5). B, D, CFP immunostaining of sectioned glabrous skin shows epidermal endings in control (B) but not mutant (D) mice, indicating a lack of peripheral terminals in Ret null nociceptors. White arrows, mature epidermal endings. White arrowheads, dermal axonal bundles. E-H, CFP immunostaining of serial DH sections from control (E&F) and mutant (G&H) mice shows sparse labeled terminals. I, Quantification of the number of CFP neurons / DRG. *n* = 14-22 DRGs from 3 animals per genotype. J, Maximal mediolateral width of sparse labeled neurons from control and mutant mice shows that the round-vs.-long distinction is still present in mutant mice. *n* = 239 (mutant), 287 (control) neurons from 3 mice per genotype. Scale bars = 100 µm (A&C), 20 µm (B&D), 50 µm (E-H).

**Figure 4.**
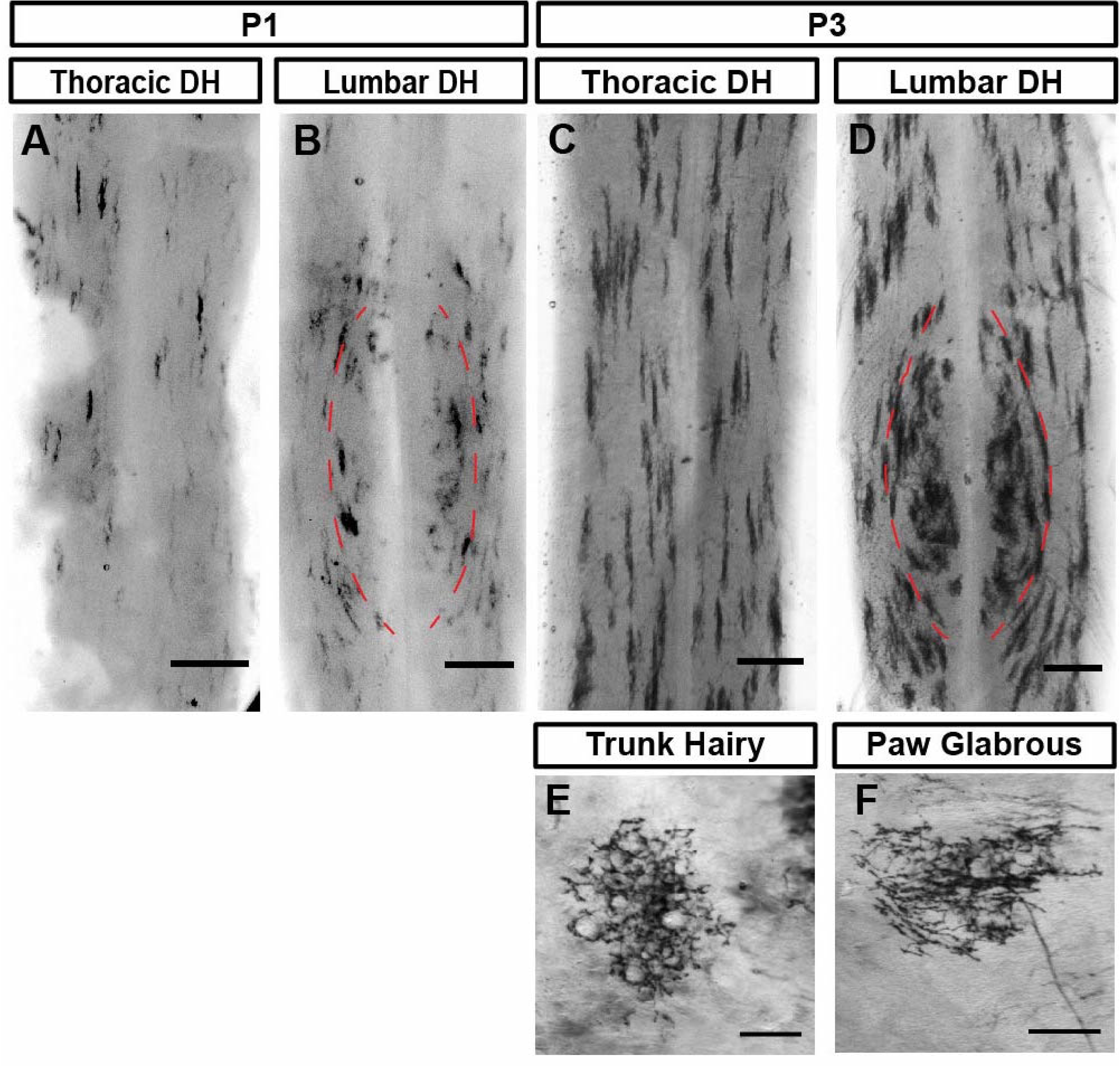
Region-specific *Mrgprd*+ DH arbors are evident during the earliest stages of innervation. A-F, AP color reaction of spinal cord (A-D) and skin (E&F) tissue from P1 or P3 *Mrgprd*^*CreERT2*^; *Rosa*^*iAP*^ mice (0.25 mg tamoxifen at E17.5). Sparse skin terminals were not seen at P1. While early postnatal nociceptors are still immature, the round vs. long distinction can be seen in B & D. Scale bars = 250 µm (A-D), 100 µm (E&F).

Though trophic factor signaling functions to control neurite arbor morphogenesis in some cell types [27, 28], here we found that deletion of the major trophic factor receptor, Ret, expressed by *Mrgprd*+ neurons [25], did not affect their DH arbor morphology. This result further indicated that disruption of the peripheral target innervation of these neurons had a negligible effect on region-specific central arbor development. Earlier work indicated that spinal cord somatotopic map formation of DRG afferents does not rely on cues from the periphery [19, 20]. Our findings expand upon this work to show that region-specific arbor morphology development also occurs independent of intact peripheral innervation.

### Sparse labeling reveals region-specific *Mrgprd*+ central arbors from the earliest stages of central innervation

Lastly, we asked whether the region-specific central terminal arbors of non-peptidergic DRG neurons might be established through postnatal reorganization (Figure 1C), or instead are present from the earliest stages of DH innervation (Figure 1D). We performed sparse genetic tracing of non-peptidergic neurons at early postnatal stages by crossing tamoxifen-dependent *Mrgprd*^*CreERT2*^ mice with *Rosa*^*iAP*^ alkaline phosphatase reporter mice (0.25 mg of tamoxifen was given at E17.5) [12]. Sparse terminals were seen after AP staining of skin and spinal cord tissue. Immature skin terminals could be seen at P3 but not P1 (Figure 4E&F), consistent with the few *Mrgprd*^*EGFPf*^ fibers in the epidermis at P1 (Figure 2). In addition, while their central arbors still appear immature at P1 and P3, region-specific arbor morphologies (the somatotopic organization of central arbors) are seen in both P1 and P3 spinal cords (Figure 4A-D). Like the mature DH organization of these afferents [12], medial lumbar enlargement (paw representation, outlined in Figure 4B, D) regions contain mediolaterally wide arbors while lateral lumbar enlargement and thoracic (proximal hindlimb, trunk) regions contain mediolaterally thin arbors (Figure 1A). It should be noted that the genetic targeting strategy utilized for this experiment may label two *Mrgprd*-lineage non-peptidergic DRG populations, one expressing *Mrgrpd* in adulthood and the other expressing *Mrgpra/b/c* genes in adulthood [12, 29]. Given that the *Mrgpra/b/c* population only represents <20% of targeted neurons [12], we believe that most if not all AP+ neurons belong to the mature Mrgprd+ population.

Taken together, while non-peptidergic central terminal arbors do show very clear postnatal layer thickness refinement (the dorsoventral axis) (Figure 2), and while early postnatal arbors have a somewhat immature morphology (Figure 4), their region-specific structure is apparent from the earliest stages of DH innervation. Earlier spinal nerve backfilling experiments indicated that DRG central projections form correct topographic innervation patterns from the earliest stages of innervation [15, 16]. Our work expands upon this to further show that region-specific central terminal arbor morphologies are also apparent shortly after the initial terminal formation (Figure 1D). This suggests that pre-patterning mechanisms may underlie the regional magnification of paw representations in the primary afferent neuropil. Future work should examine whether DRG neuron cell-intrinsic mechanisms and/or DH-intrinsic mechanisms establish the somatotopic organization of DH sensory circuits.

## MATERIALS AND METHODS

### Mouse strains

Mice were raised in a barrier facility in Hill Pavilion, University of Pennsylvania. All procedures were conducted according to animal protocols approved by Institutional Animal Care and Use Committee (IACUC) of the University of Pennsylvania and National Institutes of Health guidelines. *Mrgprd*^*EGFPf*^, *Mrgprd*^*CreERT2*^, *Rosa*^*iAP*^, and *Ret*^*f(CFP)*^ mice have been previously described [12, 21, 26, 30]. *Ret*^*null*^ allele mice were generated by crossing a conditional Ret line (*Ret*^*f/f*^) [25] with a germline Cre mouse line (*Sox2*^*Cre*^) [31].

### Genetic labeling of Mrgprd^+^ nociceptors

To sparsely label Mrgprd^+^ nociceptors, we set up timed pregnancy matings of *Mrgprd*^*CreERT2*^ mice with *Rosa*^*iAP*^ or *Ret*^*f(CFP)*^ mice. Population-level labeling was achieved through either prenatal or postnatal tamoxifen treatment. For prenatal treatment, pregnant females were given tamoxifen (Sigma, T5648) along with estradiol (Sigma, E8875, at a 1:1000 mass estradiol: mass tamoxifen ratio) and progesterone (Sigma, P3972, at a 1:2 mass progesterone: mass tamoxifen ratio) in sunflower seed oil via oral gavage at E16.5-E17.5, when *Mrgprd* is highly expressed in mouse non-peptidergic nociceptors [22].

### Tissue preparation and histology

Procedures were conducted as previously described [32, 33]. Briefly, mice were euthanized with CO2 and transcardially perfused with 4% PFA/PBS, and dissected tissue (skin, spinal cord, DRG) was post-fixed for 2 hr in 4% PFA/PBS at 4° C. Tissue used for section immunostaining was cryo-protected in 30% sucrose/PBS (4% overnight). Frozen glabrous skin and DRG/spinal cord sections (20-30 μm) were cut on a Leica CM1950 cryostat. Immunostaining was performed as described previously. DRGs for whole mount immunostaining were treated as described directly after post-fixation. The following antibodies were used: chicken anti-GFP (Aves, GFP-1020), rabbit anti-GFP (Invitrogen, A-11122). Tissue (skin or spinal cord with attached DRGs) for whole mount AP color reaction with BCIP/NBT substrate was treated as previously described. Following AP color reaction labeling, tissue was cleared in 1:2 (v:v) benzyl alcohol + benzyl benzoate (BABB) for imaging [32].

### Image acquisition and data analysis

Images were acquired either on a Leica DM5000B microscope (brightfield with a Leica DFC 295 camera and fluorescent with a Leica 345 FX camera), on a Lecia SP5II confocal microscope (fluorescent), or on a Leica M205 C stereoscope with a Leica DFC 450 C camera (brightfield). Image quantification was performed in ImageJ. Graphs and statistical analyses were created in GraphPad Prism5.

For growth normalization of skin terminal densities (Figure 2), plantar paws (n=3 animals for each age) were imaged and fitted ellipses were drawn over the six mouse foot pads. Major axis lengths (proximodistal axis) were averaged across animals. Absolute skin terminal densities were multiplied by a normalization factor: Growth-normalized density (Age) = Absolute density (Age) X (Mean paw length (Age) / Mean paw length (3 pw)).

For growth normalization of DH layer thickness (Figure 2), the maximum mediolateral width of the *Mrgprd*^*EGFPf*^ innervation layer was measured (n=3 sections from separate animals for each age). Absolute layer thickness measurements were multiplied by a normalization factor: Growth-normalized thickness (Age) = Absolute thickness (Age) X (Mean DH width (Age) / Mean DH width (3 pw)).

For single-cell width measurements in sectioned DH tissue (Figure 3), serial DH sections were imaged, and individual arbors were identified by comparing adjacent sections. The DH of lumbar enlargement sections (L3-L5) were divided into thirds based on the maximum mediolateral width of the DH. Cells with most of their width lying in the medial third were classified as “Medial Lumbar”, cells were otherwise classified as “Lateral Lumbar”.

## ACKNOWLEDGEMENTS

We would like to thank Drs. Steve Scherer, Michael Granato, and Greg Bashaw for suggestions regarding the experimental design for this study. This work was supported by National Institutes of Health (NIH) (NS083702 and NS094224 to W.L and NS092297 to W.O.) and the Klingenstein-Simons Fellowship Award in the Neurosciences to W.L.

